# Coupling between slow-waves and sharp-wave ripples organizes distributed neural activity during sleep in humans

**DOI:** 10.1101/2020.05.24.113480

**Authors:** Ivan Skelin, Haoxin Zhang, Jie Zheng, Shiting Ma, Bryce A. Mander, Olivia Kim Mcmanus, Sumeet Vadera, Robert T. Knight, Bruce L. McNaughton, Jack J. Lin

## Abstract

Hippocampal-dependent memory consolidation during sleep is hypothesized to depend on the synchronization of distributed neuronal ensembles, organized by the hippocampal sharp-wave ripples (SWRs, 80-150 Hz) and subcortical/cortical slow-waves (0.5-4 Hz). However, the precise role of SWR-slow-wave interactions in synchronizing subcortical/cortical neuronal activity is unclear. Here, we leverage intracranial electrophysiological recordings from the human hippocampus, amygdala, temporal and frontal cortices, to examine activity modulation and cross-regional coordination during SWRs. Hippocampal SWRs are associated with widespread modulation of high frequency activity (HFA; 70-200 Hz) a measure of local neuronal activation. This peri-SWR HFA modulation is predicted by the coupling between hippocampal SWRs and local subcortical/cortical slow-waves. Finally, local cortical slow-wave phase offsets during SWRs predicted functional connectivity between the frontal and temporal cortex. These findings suggest a selection mechanism wherein hippocampal SWR and cortical slow-wave synchronization governs the transient engagement of distributed neuronal populations supporting hippocampal-dependent memory consolidation.

## Introduction

Memory consolidation involves the transformation of newly encoded representations into long-term memory^1–3^. During non-rapid eye movement (NREM) sleep, hippocampal representations of recent experiences are reactivated^4,5^, along with transient synchronization of distributed subcortical and cortical neuronal populations^6–8^. It is hypothesized that the oscillatory synchrony facilitates connections between the neuronal ensembles, stabilizing memory representations^9,10^. The selection and synchronization of distant neuronal populations that participate in hippocampal-dependent memory consolidation are proposed to depend on the interaction between hippocampal sharp-wave ripples (SWRs; 80-150 Hz) and traveling subcortical/cortical slow-waves (0.5-4 Hz), but the underlying mechanisms subserving this network engagement are unclear. Here we investigated how hippocampal SWRs and subcortical/cortical slow-waves coordinate distributed neuronal populations during memory consolidation in NREM sleep.

Hippocampal SWRs are transient local field potential (LFP) oscillations (20-100 msec; 80-150 Hz in humans) implicated in planning, memory retrieval, and memory consolidation^11^. Several lines of evidence highlight the role of SWRs in sleep-dependent memory consolidation. First, memory reactivation in the hippocampus, cortical and subcortical structures peaks during SWRs^4–7,12,13^. Second, hipoccampal-subcortical/cortical functional connectivity, the prerequisite for binding of anatomically distributed reactivated memory traces is enhanced around SWRs^7,14–16^. Finally, SWR suppression or prolongation interferes with, while prolongation of SWR duration improves hippocampal-dependent memory consolidation^17,18^.

While research converges on the notion that SWR output modulates neuronal activity across brain regions during NREM sleep,SWR events are temporally biased by phases of slow-wave oscillations^19,20^. Slow-waves are present in cortical and subcortical structures^21,22^, originate in frontal areas and traverse in an orderly succession to temporal lobes and subcortical structures, including the hippocampus and the amygdala^19,22–24^. Indeed, slow-wave synchrony increases following learning^25^, and the reduction of slow-wave synchrony is correlated with memory impairment^26^. Finally, although slow-waves are ubiquitous, individual slow-wave trajectories are usually limited to a subset of cortical/subcortical areas, with ~80% of these events detected in less than half of recorded locations in humans^22^. Therefore, each SWR-associated slow-wave event could recruit and index a unique sequence of cortical and subcortical populations.

In this study, we used the broadband high frequency activity (HFA, 70-200 Hz)^27,28^ recorded from human intracranial electrodes as a metric of subcortical/cortical activity. HFA is an indirect measure of neuronal spiking from the population surrounding the electrode contact, estimated in the range of several hundred thousand neurons^29^. Consistent with the hypothesized role of SWR in synchronizing distributed memory traces, we found HFA power modulation during hippocampal SWR events in ~30% of extrahippocampal recording sites. Given the critical role of slow-waves in facilitating hippocampal-dependent memory consolidation^15^ and their confinement to local regions^22^, we hypothesizes that slow-waves organize hippocampal - cortical and cortical - cortical interactions during SWR events. Indeed, we found a strong association between SWR phase locking to cortical slow-waves and HFA modulation in the same recording site. These findings suggest that coupling of SWRs and slow-waves drive the selection of cortical populations to participate in hippocampal - cortical communication. Theoretical constructs of memory consolidation further predict transient synchronization of neuronal populations in distant cortical regions, temporally linking cortical - cortical functional connections. In support of the cooperative role of SWR and slow-waves in orchestrating cortical - cortical communication, we found that slow-wave phase alignments between two distant cortical sites predicted their neuronal population synchronization manifested by temporal HFA power correlations. These results imply a recruitment mechanism by which coupling of slow-waves and SWRs provide communication windows for long-range interactions between distributed neuronal populations, critical for hippocampal-dependent memory consolidation.

## Results

### Sleep staging and SWR detection

We recorded overnight sleep local field potentials (LFPs) in 12 subjects (573 ±18 minutes, range 480-725) simultaneously from the frontal lobe (including the orbitofrontal, medial prefrontal, dorsomedial and cingulate cortices), temporal lobe (including the insula, entorhinal, parahippocampal, inferior, medial and superior temporal cortices),amygdala (including the basolateral, lateral and centromedial amygdala), and hippocampus (Figure 1A-C). The localization of the depth electrodes was determined based on co-registered pre- and post-implantation magnetic resonance imaging (MRI), as well as registration to a high-resolution atlas. A trained researcher (B.A.M.) performed sleep staging guided by standard criteria^30^ (Figure 1D) using polysomnography (PSG) data collected from surface electrodes (i.e., electroencephalography, electrooculography, and electromyography). On average, subjects spent 287 ± 44 minutes (range 115 - 405 minutes) in NREM sleep, which represented 49.9 ± 4.1% of overnight sleep recording durations. We used depth electrodes implanted in the hippocampus to detect SWR events (see methods, Figure S1A). Hippocampal LFPs were bandpass-filtered in the SWR frequency range (80-150 Hz), rectified, and transformed to z-scores (Figure 1E). Events that exceeded five standard deviations from the mean amplitude and were beyond a one-second window of the nearest interictal epileptic discharges (Figures S1A-B) were classified as SWRs. As shown in Figure 1E, the morphology of grand-average SWR (n = 12 subjects; 44965 SWRs) and the numbers of SWR events per hippocampal electrode (1653 ± 274) were consistent across subjects and in line with previous reports from humans and non-human primates^20,31–34^

**Fig. 1.**
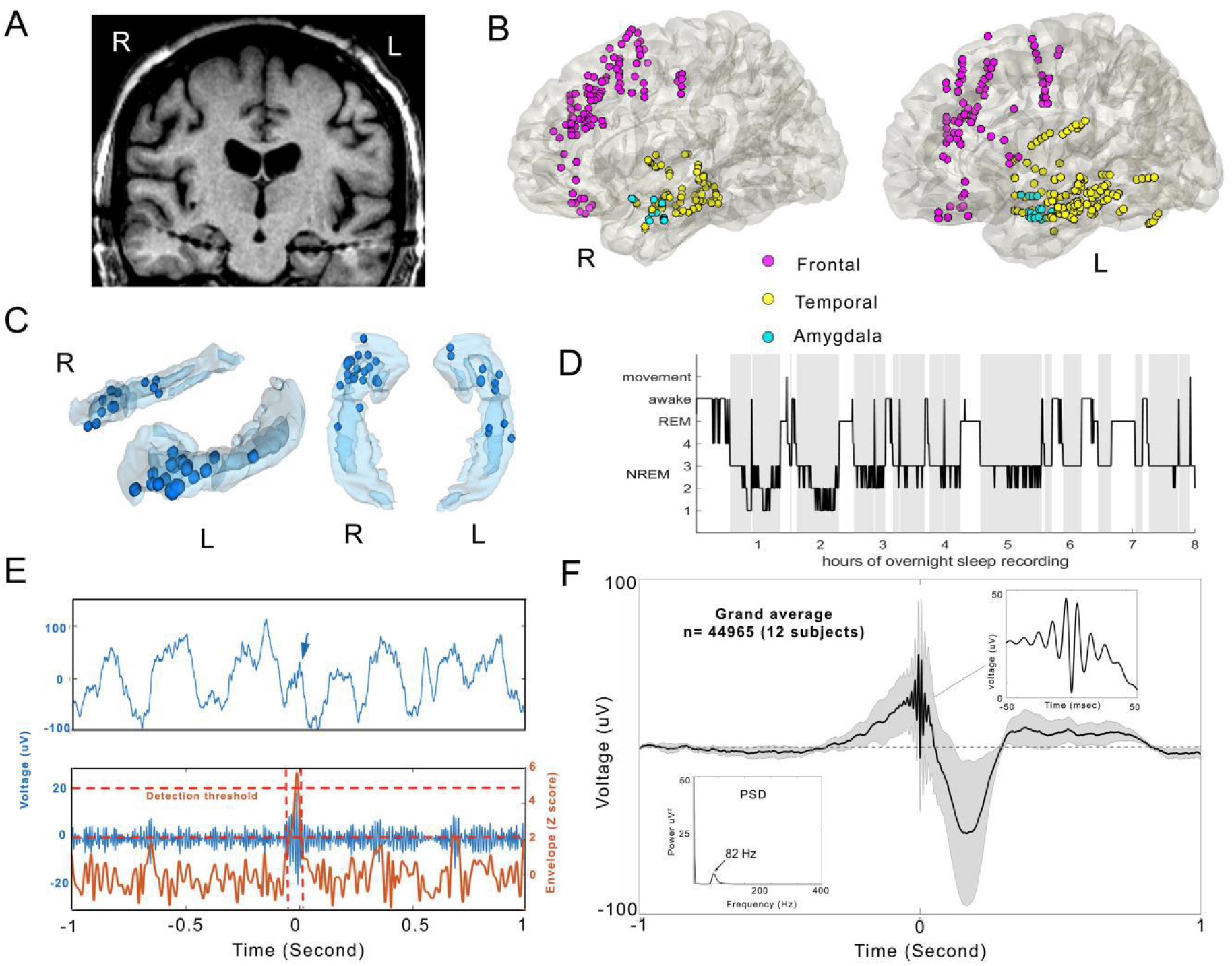
Recording locations, sleep staging, and sharp-wave ripple detection. **a,** Coronal MRI image from an example subject, showing the depth electrodes targeting the hippocampus bilaterally. Thicker areas along the electrode shafts represent the individual recording contacts. **b**, Anatomical distribution of extra-hippocampal recording locations (amygdala - cyan, n = 36; temporal cortex - yellow, n = 180; frontal cortex - magenta, n = 189; medial views of the right (R) and left (L) hemispheres. Regional electrode distributions across individual subjects are shown in the Supplementary Table 1. **c**, Hippocampal electrode localizations (n = 33) from all the subjects, shown on a 3-dimensional hippocampus model. Blue dots indicate the individual electrode locations in the left (L) or right (R) hippocampus. **d**, Overnight sleep hypnogram from an example dataset. The non-rapid eye movement (NREM) stages 2-4 were used in the analysis (grey).**e**, Example of detected sharp-wave ripple (SWR) and illustration of the detection algorithm. Top: raw LFP trace around the SWR (arrow). Bottom: blue - the LFP trace shown in the top plot, band-pass filtered in the SWR range (80-150 Hz). Orange - z-scored envelope of the filtered trace. SWR detection was based on the coincidence of two criteria: 1) SWR envelope peak crossing the mean + 5SD (upper threshold, top dashed orange line) and 2) SWR envelope around the candidate upper threshold crossing exceeding mean + 2SD for 20-100 msec (lower threshold, bottom dashed orange line). **f**, Grand-average SWR-centered raw LFP (mean ± SEM; n = 12 subjects). Time 0 corresponds to SWR peak. Top right inset: Several oscillatory cycles around the grand-average SWR peak reflect the oscillatory nature of detected SWR events, lasting several tens of msec. Bottom left inset: Power spectral density (PSD) averaged across all detected SWRs. Unimodal peak (82 Hz) suggests the lack of contamination with epileptic activity, which is typically reflected as an additional PSD peak in > 200 Hz range^33^.

### Hippocampal sharp-wave ripples modulate subcortical and cortical high frequency activity

Functional MRI studies show a widespread peri-SWR activity modulation^35^, but the precise timing and anatomical distribution of the neuronal activity is unclear. Therefore, we leveraged millisecond temporal resolutions and broad anatomical coverage of intracranial electrophysiological recordings to measure HFA power (a proxy of neuronal population activity) during SWR windows (± 250 msec, relative to SWR peak). We paired each hippocampal recording site containing a minimum of 100 SWRs/overnight recording session with simultaneously recorded extra-hippocampal recording sites and operationally defined the pairs as target sites (n = 1308, 625 ipsilateral and 683 contralateral to SWR location; Figure 2A). All of the electrodes used in the analysis were localized in gray matter (see Methods). Based on the presence or absence of significant peri-SWR HFA modulation (see Methods), target sites were classified as either HFA+ or HFA-. We found significant peri-SWR HFA power modulation in 28.1% (368/1308) of target sites. Linear regression analysis showed significant main effects of region (F(1,1304) = 219.6, p < 10^-10^) and hemisphere (F(1,1304) = 133.5, p < 10^-10^) on the percentage of target sites showing peri-SWR HFA modulation, as well as a region by hemisphere interaction (F(1,1304) = 109.5, p < 10^-10^). Moreover, the percentage of peri-SWR HFA modulated subcortical/cortical target sites was significantly higher when SWRs originate from the same hemisphere (278/625, 44.5%) versus SWRs arising from the contralateral hemisphere (110/683; 16.1%; Figure S2A; chi-square: χ^2^ (1, n = 1308) = 116.2, p < 10^-10^). The highest percentage of modulated sites were in the amygdala (88.6% ipsilateral and 30.2% contralateral to SWR), followed by the temporal (67.6% ipsilateral and 19.2% contralateral) and the frontal cortex (15.1% ipsilateral and 12.2% contralateral; Figure 2C, S2B). Detailed statistical comparisons of regional and hemispheric peri-SWR HFA modulation are shown in Supplementary Tables 2A and B. We classified peri-SWR HFA power modulations as either: 1) positive (increased HFA power); 2) negative (decreased HFA power); or 3) mixed (both periods of increased and decreased HFA power; Figure 2B). Positive modulations were the most common modulation class (266/368, 72.3% of HFA-modulated target sites), followed by mixed-modulations (74/368, 20.1%) and negative-modulations (28/368, 7.6%). Overall, these findings suggest an anatomically - specific engagement of neuronal populations during SWR windows.

**Fig. 2.**
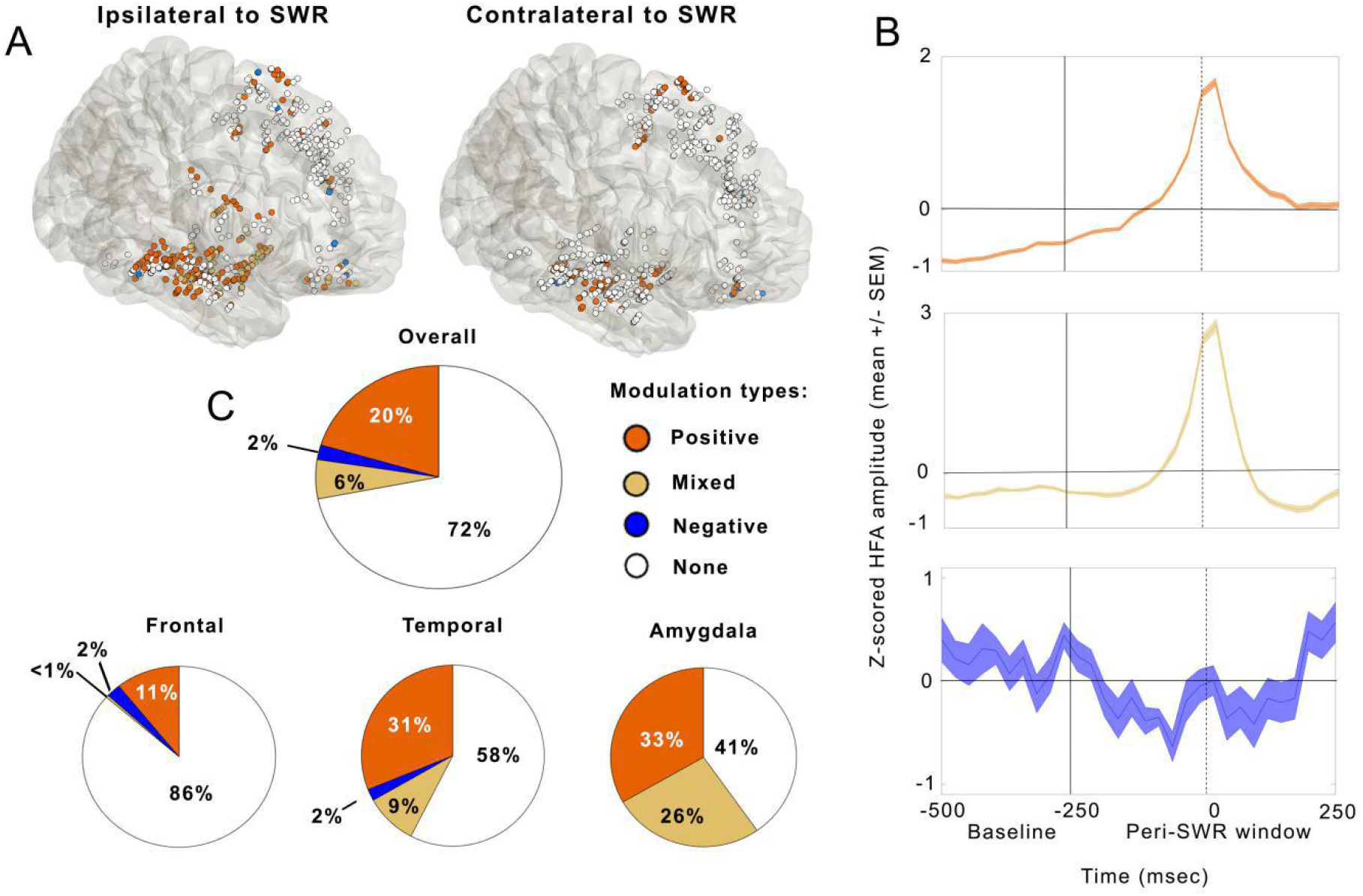
High frequency activity (HFA) modulation around sharp-wave ripples (SWRs). **a**, Anatomical distributions of peri-SWR HFA modulation ipsilateral (left) and contralateral (right) to SWR location. Orange - positive-modulation, blue - negative-modulation, ocher - mixed modulation and white - no modulation (1308 target sites, 625 ipsilateral and 683 contralateral to SWR location). Target sites are defined as extra-hippocampal recording locations. When SWRs are recorded from multiple hippocampal channels in the same subject, locations of target sites are minimally jittered for visualization purposes. The most common significant peri-SWR HFA modulations are positive-modulations ipsilateral to SWR location in temporal and amygdala sites. See also Movie S1 for the temporal dynamics of peri-SWR HFA modulation. **b**, Average z-scored peri-SWR HFA time-courses (mean ± SEM) from all the target sites showing a given modulation type (positive-modulation - top, mixed modulation - middle, negative-modulation - bottom). Tick vertical line denotes the boundary between baseline (−500 to −250 msec, relative to SWR peak time) and peri-SWR window (± 250 msec around SWR peak time). SWR peak time is denoted by the vertical dashed vertical line. **c**, Percentages of significant peri-SWR HFA modulation overall (top chart) and at the regional level (bottom charts). Orange - positive modulation, blue - negative modulation, ocher mixed modulation and white - no modulation.

While the previous analysis characterizes modulation at the level of individual target sites, memory consolidation theories predict coordinated activity of distributed neuronal populations around hippocampal SWR windows. Thus, we hypothesized that these temporal profiles co-vary across target sites around the time of SWR. To examine the low dimensional representation of neural population activity, we performed a principal component analysis (PCA) on the HFA time-courses derived from a pseudo-population of HFA modulated target sites, ipsi- and contra-lateral to hippocampal SWRs (n = 368). Consistent with the prediction of a low dimensional space of peri-SWR HFA dynamics, the first three principal components (PCs) accounted for 55.5%, 15.0%, and 6.2% of the explained variance (EV), respectively (cumulative EV = 76.7%; Figure 3A). Reconstructed activity time-courses of these three PCs revealed distinct dynamics, with PC1 activity showing a symmetric increase around the hippocampal SWR, while PC2 time-course showing a persistent activity increase following the SWR (Figure 3B). The PC3 time-course is bimodal, characterized by activity peaks both before and after the SWR peak (Figure 3B). For each of the first three PC scores, a two-way ANOVA (with region and hemisphere relative to SWR location as main factors) showed significant main effects of region (F’s(2,367) > 7, p’s < 10^-3^) and hemisphere (F’s(1,367) < 8, p’s < 10^-5^), as well as region by hemisphere interactions (F’s(1,2) >11, p’s < 10^-4^), except for PC3 (F(2,367) = 2.37, p = 0.1). Remarkably, PC distributions showed regional specificity, with PC1 dynamics represented in the amygdala and the temporal cortex, while PC2 dynamics characterized the peri-SWR HFA in the frontal cortex (Figures 3C). Detailed statistical comparisons of regional PC weights are shown in Supplementary Table 3. Further, for different brain regions, the state-space trajectories around the time of the SWR show distinct dynamics (Figure 3D). The trajectories derived from the amygdala and the temporal cortex peri-SWR HFA are characterized by increased velocity around the SWR peak and returning thereafter to the baseline (Figure 3D). In contrast, the state space trajectory derived from the frontal cortex peri-SWR HFA is characterized by a slower velocity along the PC2 axis, without returning to the baseline within the 250 msec after the SWR peak. This analysis reveals low-dimensional peri-SWR HFA dynamics, which could be organized by oscillatory synchrony. Based on the role of slow-waves in long-range synchronization supporting the hippocampal-dependent memory consolidation^15,20,25,36^, we next sought to discern the potential role of SWR and slow-wave interactions in facilitating cortical - cortical communication.

**Fig. 3.**
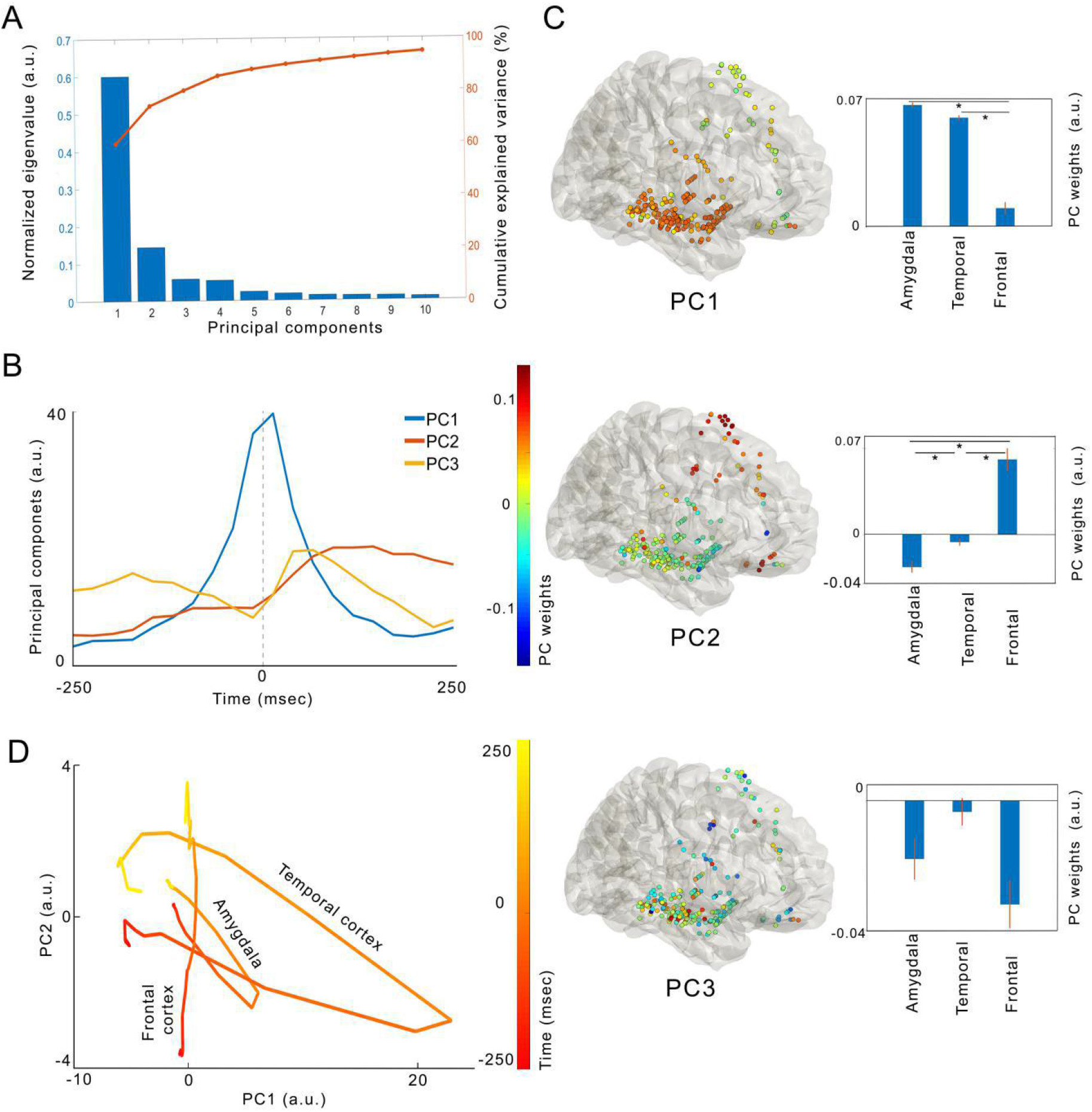
Principal component analysis shows low-dimensional and regionally-specific peri-SWR HFA dynamics patterns. **a**, Blue bars - normalized eigenvalues and orange line - cumulative explained variance (EV) for the first ten principal components. The first three principal components show significance (see Methods), with cumulative EV ~80%. **b**, Reconstructed peri-SWR HFA time-courses (± 250 ms) of the first three principal components, centered on the SWR peak (dashed gray vertical line). PC1 activity is characterized by a symmetric peak around SWR, while the PC2 activity shows a protracted increase following SWR. PC3 activity time-course is bimodal, with distinct peaks bot before and after the SWR peak. **c**, Left column: Glass brain maps of different PC weights in the hemisphere ipsilateral to SWR peak, denoted by color. Right column: the first three PC weights. Right column: Pseudo-population regional means of the PC weights for the corresponding PCs from the left column. PC1 weights in amygdala and temporal cortex are higher, relative to frontal cortex, while the anatomical distribution of PC2 weights shows the opposite pattern (two-tailed t-test, p < 0.05, statistical details in Supplementary Table 3). Data is shown as mean ± SEM. **d**, Projections in state space of the ipsilateral regional average trajectories for the first two PCs, which were showing regional weight differences. Line color gradient denotes the time domain, ranging from −250 msec to 250 msec, relative to the SWR peak. Note the increased state-space distances between adjacent time-points closer to SWR peak time, reflecting the higher velocity in state space. Amygdala and temporal cortex trajectories tend to move along both PC1 and PC2 axes, while the frontal cortex trajectory moves mostly along the PC2 axis. In addition, the amygdala and temporal cortex trajectories return close to origin by the end of the peri-SWR period. In contrast, the frontal cortex trajectory remains in a different part of state space, suggesting prolonged activity in frontal populations modulated around SWR.

### SWR synchrony with subcortical/cortical slow-waves predicts local activity modulation

If slow-waves rhythmically modulate neuronal excitability^21,22^, we hypothesized that the synchrony between hippocampal SWRs and slow-waves at individual subcortical/cortical target sites would predict local HFA modulation. For each target site, we define SWR-slow-wave synchrony as significant phase-locking between hippocampal SWRs and local slow-waves (Rayleigh test, p < 0.05, with Benjamini-Hochberg correction for multiple comparisons; Figure 4A and Methods). First, we found a higher percentage of target sites with SWR-slow-wave synchrony in the ipsilateral, compared to contralateral hemisphere, relative to the SWR location (39.4% (246/625) vs. 13.8% (94/683); χ^2^ (1, n = 1308 target sites) = 131.3, p < 10^-10^; Figure 4B and S4A). Second, the percentage of SWR-slow-wave synchrony was significantly higher for HFA+, relative to HFA- sites (ipsilateral HFA+ = 71.7% (185/258), HFA- = 16.6% (61/367), χ^2^ (1, n = 625 target sites) = 182.5, p < 10^-10^; contralateral HFA+ = 37.3% (41/110), HFA- = 9.3% (53/573), χ^2^ (1, n = 683 target sites) = 53.4, p < 10^-10^), Figure S4B). Further, a higher percentage of SWR-slow-wave synchrony in HFA+ relative to HFA- target sites was present in all recorded brain regions except in the ipsilateral amygdala (89.8% HFA+ (35/39) or 60% (3/5) HFA- sites (χ^2^(1, n = 44 recorded sites) = 3.33, p = 0.07; Figure S4C). In all other recorded brain regions, including the contralateral amygdala, the percentage of SWR-slow-wave synchrony was 2-4 times higher in HFA+, relative to HFA-target sites (Figure S4C; Supplementary Table 4, all p’s < 10^-3^).

**Fig.4.**
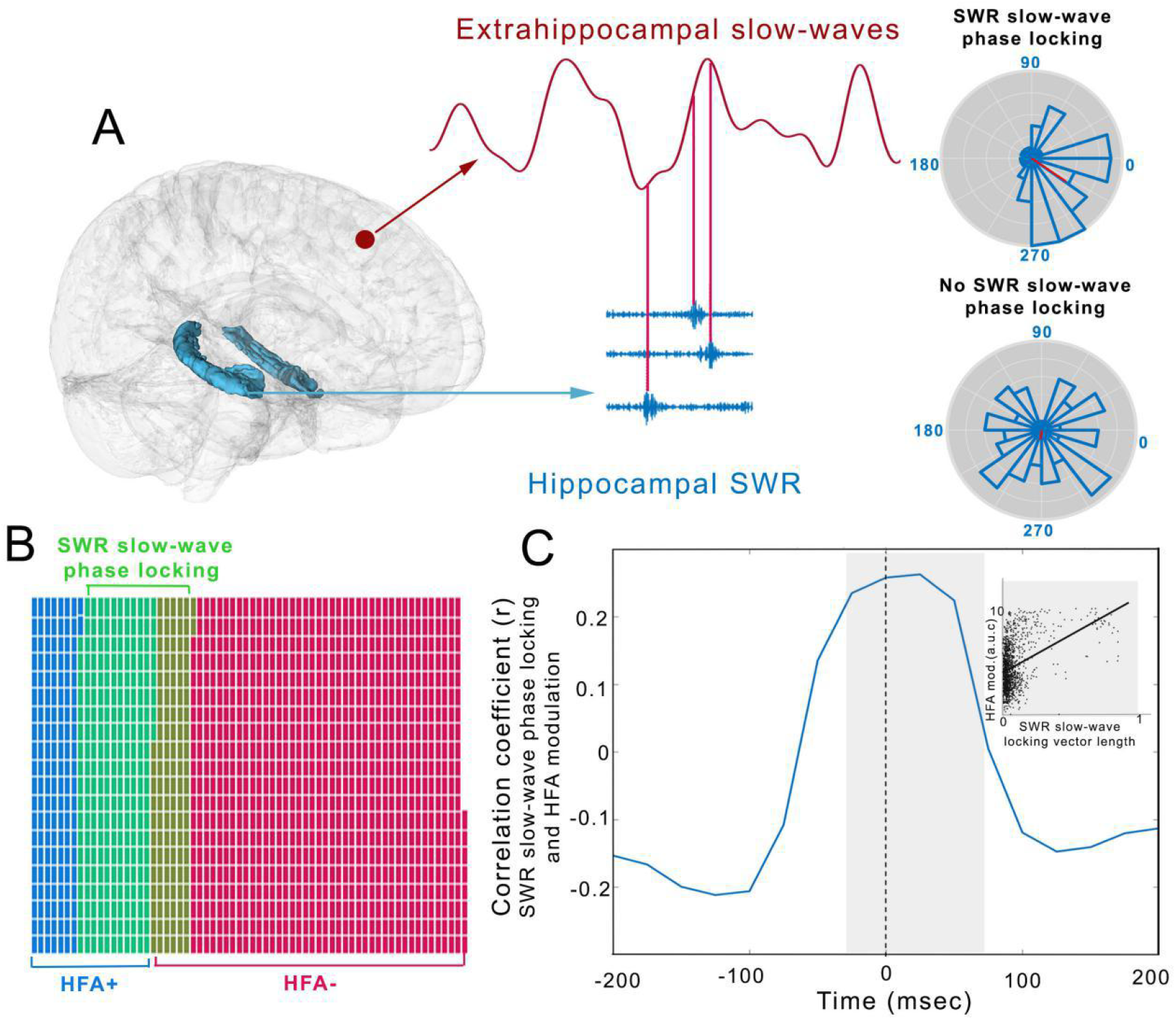
SWR synchrony with subcortical/cortical slow-waves predicts local peri-SWR HFA modulation. **a,** Illustration of the SWR-slow-wave phase locking. Local slow-wave phases from an example extra-hippocampal target site (maroon) corresponding to SWR peak times on simultaneously recorded hippocampal channel (blue) are used to construct the target site-specific circular distribution of slow-wave phases coinciding with hippocampal SWR peaks. The distribution uniformity is tested with the Rayleigh test (p < 0.05, Benjamini-Hochberg correction for multiple comparisons). Based on the presence of significant non-uniformity, the target sites are classified as showing SWR-slow-wave synchrony (SWR-slow-wave+) or no synchrony (SWR-slow wave-). Right column: Polar plots showing the distribution of slow-wave phases at SWR peak times for the SWR-slow wave+ (top) and SWR-slow-wave- (bottom) example target sites. **b**, Venn diagram showing the larger overlap between the target sites showing SWR-slow wave synchrony (green) and HFA+ (blue), relative to HFA- (red) target sites (χ2(1, n = 1308) = 333.9, p < 10^-10^). Individual target site are represented as squares. **c**, Correlation between the HFA modulation strength and extent of SWR-slow wave phase locking (as denoted by the length of r vector), calculated for 100 msec sliding windows (25 msec step size). The peak correlation is shown for the window ranging from −25 to 75 msec around the SWR peak (Spearman r = 0.26, p < 10^-10^). Inset: scatter plot illustrating the positive correlation between vector length and HFA modulation strength in the highlighted window. For both plots, n = 1308 target sites.

After establishing association between the presence of SWR-slow-wave synchrony and local HFA modulation, we tested the relation between the SWR-slow-wave synchrony magnitude and the strength of local peri-SWR HFA modulation. We define the synchrony magnitude as the length of mean vector r (Rayleigh test) and peri-SWR HFA modulation strength as the sum of HFA power (baseline-corrected and z-scored). SWR-slow-wave synchrony was positively correlated with HFA modulation strength around the SWR peak (Figure 4C), reaching the maximum correlation at 25 msec (Spearman r = 0.26, p < 10^-10^; n = 1308 target sites).

The anatomical selectivity of SWR-slow-wave synchrony is not attributable to slow-wave power, as this parameter is not different between target sites with and without significant SWR-slow-wave synchrony (Figure S4D; independent samples two-tailed t-test, t(1306) = 0.19, p = 0.85; n = 1308). In summary, these findings indicate that the synchrony between hippocampal SWRs and target site slow-waves is a potential mechanism for selecting neuronal populations to participate in widespread synchronous activity during SWRs.

### Slow-wave phase difference during sharp-wave ripples predicts cortico-cortical coupling

As demonstrated by simultaneous recordings from multiple locations in the monkey visual cortex, neuronal population locking to specific phases of local gamma oscillations determines their functional coupling^37^. Similarly, we hypothesized that the HFA locking to slow-wave phases during SWR windows determines the strength of pairwise coupling between populations across the subcortical/cortical regions, organizing functionally connected transient modules^36^. HFA coupling is defined as the temporal correlation between the HFA analytic amplitude time-courses at different target sites. To test this hypothesis, we first quantified the target site pairwise HFA coupling during SWR windows, using a generalized linear model^38^. This method allows pairwise coupling estimation while factoring out the contribution of common coupling to global neuronal activity. For a given target site pair (A and B), peri-SWR HFA time-course on target site A (HFA-A) was modeled based on two regressors, peri-SWR time-course on target site B (HFA-B) and average population peri-SWR HFA time-course (HFA-pop). We define the population activity as HFA from all the simultaneously recorded target sites, except A and B. This method produced a pair-specific coupling coefficient (β_0_), with amplitude reflecting the coupling strength and sign reflecting positive or negative coupling. We repeated this analysis on the same target site pair, but with HFA-A and HFA-pop used as regressors for modeling HFA-B. Next, we tested the association between the pairwise HFA coupling and local slow-wave phases at the time of SWR. The analysis focused on a subset of target site pairs meeting both of the following criteria: 1) HFA modulation (HFA+; Figure 2) during the SWR window and 2) SWR synchrony with local slow-waves (Figure 4). For each target site pair, we correlated the coupling coefficient β0 and the slow-wave phase difference (ϕ_diff_) between two target sites at the SWR occurrence time (Figure 5A-C). When analyzing target site pairs from all the regions in both hemispheres together, β0-φdiff correlation was not significant after correcting for multiple comparisons using the Benjamini and Hochberg method (Spearman correlation, n = 1716 pairs, r = −0.04; p = 0.045). These findings suggest that slow-wave phase differences did not predict HFA coupling at the global level. We next computed the β_0_-φ_diff_ correlation for within-region (e.g., amygdala - amygdala pairs) and cross-region sets (e.g., frontal - temporal cortex pairs) with at least 20 simultaneously recorded target site pairs (Figure 5A). The frontal-temporal target site pairs ipsilateral to the side of hippocampal SWR showed significantly negative β_0_-φ_diff_ correlation (Figure 5D; Spearman correlation, n = 166 pairs; r = −0.26, p < 0.001). Negative β_0_-φ_diff_ correlation reflects the positive functional coupling with small slow-wave phase offsets at the time of SWR and negative coupling with larger phase offsets. These results show that the local slow-wave phase difference at the time of SWR occurrence determines the HFA coupling between the frontal and temporal target sites. This coordination mechanism appears specific for frontal-temporal interactions as there was no significant β_0_-φ_diff_ correlation within or across any other regions (see Supplementary Table 5 for statistics).

**Fig. 5.**
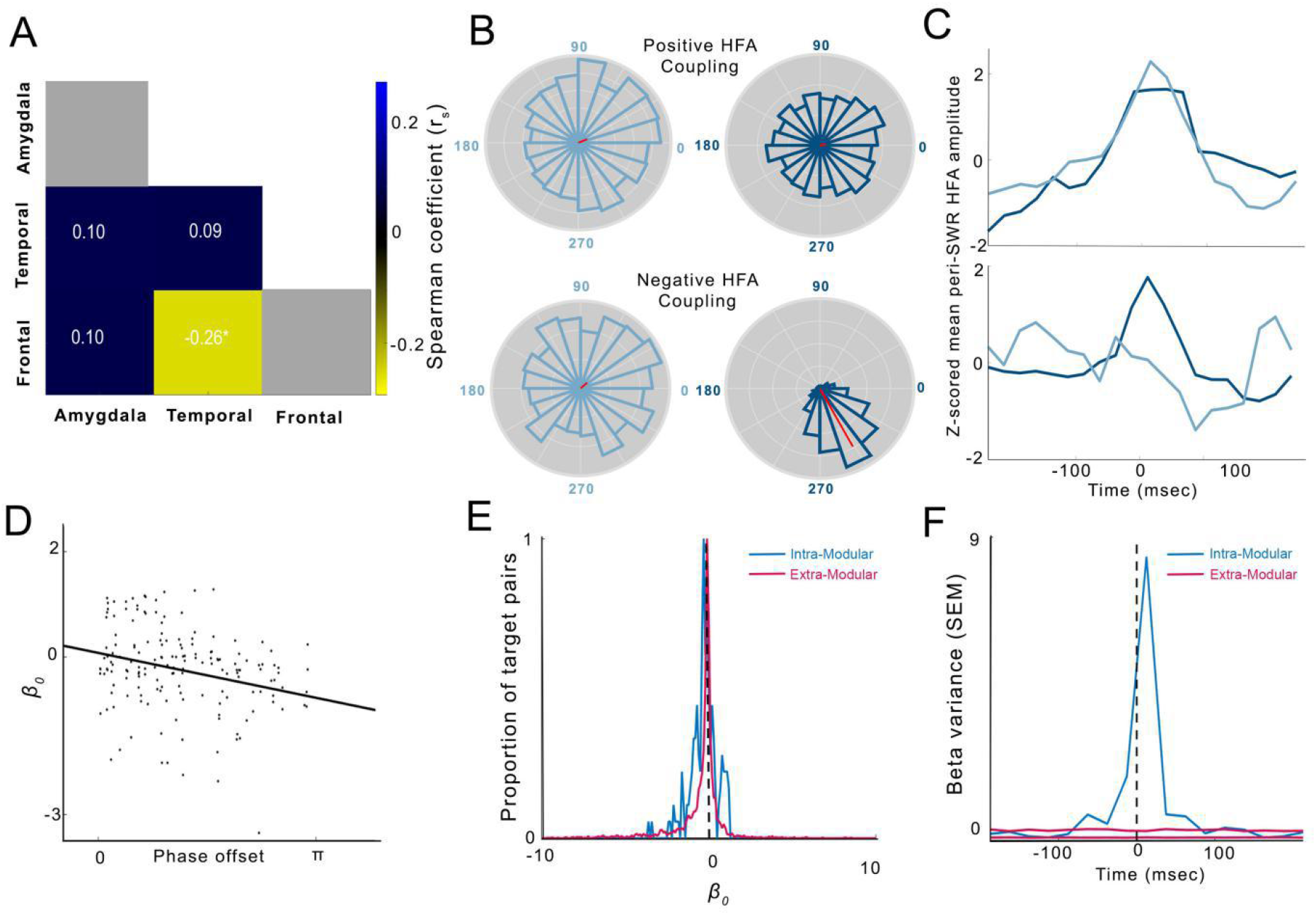
Correlations between the pairwise SWR-slow-wave phase difference (φ_diff_) and HFA coupling (β_0_). **a,** Correlation matrix for ipsilateral within- and cross-structure SWR-slow wave phase-locking difference φ_diff_ and HFA β_0_ Color denotes the Spearman r value for a given region pair. White color denotes the region pairs with < 20 simultaneously recorded target site pairs, which were excluded from the analysis.**b**, Examples of SWR-slow wave phase distributions for the target site pairs showing highly positive (top row) or negative (bottom row) HFA coupling (β0). **c**, Peri-SWR HFA time courses from the target site pairs in **b**. Positive or negative HFA coupling is associated with small or large φ_diff_, respectively. **d**, Scatter plot showing the negative correlation between the φ_diff_ and HFA β_0_ forthe frontal-temporal target sites from the hemisphere ispsilateral to SWR location (Spearman correlation (r = −0.26, p < 0.001; n = 166 pairs). Negative correlation denotes the functional coupling between the pair of frontal-temporal sites becoming positive with decreased local slow-wave phase differences at the time of SWR. **e**, Histogram of pairwise HFA β_0_ coefficients for the intra-modular (blue) and extra-modular (maroon) frontal-temporal target site pairs. Distribution of intra-modular β_0_ is significantly wider dispersed around zero, denoting the higher proportion of strongly positively or negatively coupled intra-modular pairs, relative to both extra-modular and combined target pairs (Ansari-Bradley dispersion test, p < 0.002). **f**, Variance of pairwise frontal-temporal intra-modular (blue) and extra-modular (maroon) β_0_ coefficients during the window around SWR. Extra-modular variance is shown as 1^th^ and 99^th^ percentile of 100 random samples for each sliding window. SWR peak time is shown as dashed line.

If slow-waves organize transient functional modules around the time of SWRs^36,39^, then the HFA+ target sites organized by slow-waves (intra-modular) would show stronger mutual coupling, relative to coupling with other target sites (extra-modular). Also, the strong intra-modular coupling should appear transiently, locked to SWR events. We define coupling strength as the dispersion width of HFA pairwise coupling coefficients (β_0_). We observed that the intra-modular frontal-temporal coupling strength was significantly stronger, relative to extra-modular (Ansari-Bradley test, n = 166 intra-modular and 3026 extra-modular target site pairs, p < 0.002; Figure 5E). These findings are conceptually similar to the wider dispersion of spike train coupling coefficients between the intra-modular, relative to extra-modular grid cell pairs, which indicated the modular organization in the entorhinal cortex^38^.

Finally, to test if the higher frontal-temporal intra-modular coupling strength is transient and locked to hippocampal SWRs, we compared the variance (expressed as the standard error of mean, SEM) of intra- and extra-modular coupling strengths using a sliding window (100 msec width, 25 msec step size), centered between −200 and 200 msec around SWR peak. To avoid the effect of unequal sample size on variance, for each time window, we randomly sampled 100 sets of extra-modular target site pairs, equal in size to the intra-modular set (n = 166). Indeed, intra-modular connectivity variance was significantly higher compared to extra-modular connectivity within ±100 msec window of hippocampal SWR (Figure 5F; >99th percentile of the extra-modular set; p < 0.01). In summary, these results confirm the presence of modular organization of distributed neural activity during SWR windows, which could support the functional segregation of populations participating in global peri-SWR activation preventing interference from random background activity^10^. Furthermore, the transient nature of modular activity locked to SWRs suggests that HFA coupling does not reflect the stable connectivity patterns during NREM sleep, but emerges from SWR-slow-wave coordination.

## Discussion

These findings reveal widespread modulation of brain activity during hippocampal SWRs during time windows of memory reactivation in the hippocampus and associated subcortical/cortical structures^8,11,40^. In addition, there is a strong correlation between peri-SWR activity modulation in a given location and synchrony between hippocampal SWRs and local subcortical/cortical slow-waves. Thus, SWR-slow-wave synchrony may act as a mechanism for selecting distributed populations recruited simultaneously with hippocampal memory traces reactivated during SWRs. Finally, functional coupling between the pairs of sites in frontotemporal network during SWR windows is correlates with phase offsets between the local slow-waves. This could underlie the formation of temporary neuronal coalitions around the SWRs, as predicted by memory consolidation theories^3,36,41^.

### Peri-SWR HFA modulation

Most of the peri-SWR HFA modulations were positive, in line with the observations of widespread increase in cortical blood oxygen level-dependent (BOLD) signal and the higher probability of cortical gamma bursts during peri-SWR windows^32,35^. Mixed peri-SWR HFA modulations, were mostly present in temporal lobe ipsilateral to SWR location, manifested as HFA increase around the SWR peak, followed by decrease 100-200 msec later (Figure 2B, Supplementary Movie 1). Peri-SWR HFA negative-modulations were of lower amplitude and most common in the frontal cortex (~15% of modulations in that region; Figure 2C). The presence of both positively and negatively modulated peri-SWR HFA in frontal areas could reflect consistent co-occurrence of local up-to-down or down-to-up state transitions with SWRs. However, the low SWR density during cortical down-states^7,22,42^ and the wider spatial synchronization of down-states^22^, would likely result in more widespread peri-SWR HFA negative modulation than observed in the present data. Another possible interpretation of frontal peri-SWR HFA negative modulations could be the recruitment of local inhibitory networks, similar to increased firing of inhibitory interneurons in deep layers of rodent prefrontal cortex during SWRs^43^.

Peri-SWR HFA modulation is region- and hemisphere-dependent, with higher percentages ipsilateral to SWR location (Figures 2C, D and S2A, Supplementary Movie 1), as well as in the temporal lobe structures (amygdala and temporal cortex), where it reached >70-80% of target sites in the ipsilateral and 20-30% in the contralateral hemisphere. Interestingly, hemispheric differences were not present in the frontal cortex, where 15-20% of sites were modulated irrespective of the SWR origin laterality (Figures 2C, E, S2B, and C). In addition to classifying the peri-SWR HFA modulation based on direction of change, we applied PCA to assess the dimensionality of peri-SWR HFA time-course. PCA revealed the low dimensionality of peri-SWR HFA time-courses, with ~80% of variance explained by the first three principal components (Figure 3A). This low dimensionality shows a robust regional mapping, with the sites carrying high PC1 weights, characterized by symmetric peri-SWR activity increase located predominantly in the amygdala and temporal cortex. On the other hand, the sites carrying high PC2 weights, characterized by persistent delayed increase following SWR - located mostly in the frontal cortex (Figure 3B). Widespread and low-dimensional peri-SWR HFA modulation suggests the synchronized activation of distributed neuronal ensembles during SWR windows, a necessary prerequisite for binding of distributed memory traces underlying hippocampal-dependent memory consolidation^10^. Higher presence of HFA modulation in temporal cortex and amygdala could reflect enhanced connectivity between the hippocampus and temporal lobe structures, relative to hippocampal-frontal connectivity, although the frontal cortex is one of the principal hippocampal target areas outside the temporal lobe^44^.

### SWR-slow-wave synchrony predicts local HFA modulation

We demonstrate that the consistent relation between the hippocampal SWR timing and subcortical/cortical slow-wave phases (SWR-slow-wave synchrony) strongly predicts a subset of local populations participating in global peri-SWR activation^35^, proposed to be involved in memory consolidation^36,40^. The sites with significant peri-SWR modulation or no modulation co-exist in the same brain structures (Figure 2A). Such a selective pattern of peri-SWR HFA modulation suggests that besides the necessary anatomical connectivity, the local subcortical/cortical slow-waves could provide a gating mechanism that enables the peri-SWR HFA modulation. The traveling slow-waves tend to emerge from frontal lobe and spread in the anterior-posterior direction^23^, but the site of origin, traveling directions and velocities across individual waves are highly variable^22,23^. Moreover, ~85% of slow-waves in the human brain show relatively limited extent, invading less than half of recorded locations^22^. Although the slow-wave trajectories are constrained by anatomical connections^45^, slow-waves are theoretically capable of sampling from a large combinatorial space of subcortical/cortical populations, enabling their synchronous activity during individual peri-SWR windows. Slow-waves are regulated by local learning history and appear with higher amplitude and more often in the regions involved in recent learning^46,47^.This could be due to locally-regulated mechanisms, such as learning-dependent changes in excitatory-inhibitory balance^48^, that creates the path of least resistance for the traveling slow-waves, biasing their trajectories towards populations modified by recent learning. Hence, the slow-wave trajectory could be biased towards visiting the areas involved in recent learning, thereby representing the dynamic selection mechanism for synchronized reactivation of distributed memory traces around the SWRs, facilitating hippocampal-dependent memory consolidation^15^.

### Slow-wave phase during SWRs determines cortico-cortical functional coupling

Slow-waves synchronization in anatomically distributed neuronal populations during peri-SWR periods could facilitate formation of transient neuronal coalitions providing a mechanism for binding of distributed memory traces^36^. We demonstrate that the strength and sign of long-distance interactions in the fronto-temporal network during SWR windows are dependent on the phase differences between local slow-waves. Critically, this correlation is present after factoring out the common coupling to global brain activity, by applying a generalized linear model (GLM). This is a widely used approach for quantifying the relative contributions of multiple predictors on the activity of single neurons^8,38,49^ and neuronal populations^50^. The relation between the oscillatory phase offset and functional connectivity strength between the different local populations has been previously demonstrated in the visual cortex^37^.

In addition, the distribution of coupling coefficients for the temporal-frontal intra-modular pairs, defined by showing peri-SWR HFA modulation and SWR-slow-wave synchrony, was much wider than coupling with other sites (extra-modular; Figure 5E). Similarly, grid phase offset-dependent strong positive or negative spike train correlations were found only for the grid cell pairs belonging to the same functional module, resulting in wider distribution of intra-modular coupling strengths^38^. In general, stronger intra-modular connectivity is a hallmark of functional organization in the brain^39^. These results suggest that slow-waves functionally segregate the subspace of anatomically distributed neuronal populations, organized by their coupling to local slow-wave phase. HFA coupling for most of the temporal-frontal target site pairs is negative suggesting a relatively high dimensional communication space occupied by a larger number of subspaces defined by the local slow-wave phase differences.

Finally, the slow-wave phase-dependent functional coupling between the cortical sites does not reflect the stable functional connectivity matrix during NREM sleep, but is temporally coupled to SWRs (Figure 5F). Such a dynamic suggests the role of slow-wave phase-organized distributed neuronal coalitions in hippocampal-dependent memory consolidation supporting binding of distributed memory traces^3,10^.

### Summary

Various models of hippocampal-dependent memory consolidation implicate binding between the hippocampal memory traces reactivated during SWRs and the subcortical/cortical populations encoding various aspects of the same experience^2, 3^. These results suggest the critical role of a consistent phase relation between the hippocampal SWRs and subcortical/cortical slow-waves for the selection of local populations active during SWR windows. In addition, the local slow-wave phases during SWRs predict the functional coupling between the distant cortical populations, enabling the plasticity necessary for binding of distributed memory traces. Our findings implicate SWR-slow-wave synchrony as a core mechanism affecting the content, fidelity and strength of consolidated memories.

## Materials and Methods

### Experimental Design

The first objective of this study was to map the spatio-temporal modulation of neural activity around the SWR windows in human brain, as reflected by the HFA amplitude, a proxy measure of population activity in proximity of electrode tip. Additional objective was to assess the dependence of local subcortical/cortical peri-SWR HFA modulation on the coupling between hippocampal SWRs and local slow-waves. Finally, we aimed to assess the relation between the local slow-wave phase differences during SWR windows and HFA functional coupling across the recorded structures, including the amygdala, temporal and frontal cortices.

Twelve pharmacoresistant epileptic patients (7 males, 5 females, age 38±4 (mean ± SEM), range 24-57) undergoing presurgical evaluation of seizure foci at the University of California Irvine (UCI) Medical Center were included in the study based on written informed consent. All the procedures were performed in accordance with the UCI Institutional Review Board. The subjects were stereotactically implanted with 6-10 intracranial depth electrodes (Integra or Ad-Tech, 8-10 macroelectrodes with 5-mm inter-electrode spacing) under robotic assistance (Rosa Surgical Robot, Medtech, New York, NY). Electrode placements were driven strictly by clinical diagnostic needs and included the unilateral or bilateral implants in the hippocampus, amygdala, temporal and frontal cortices. Details of individual patient electrode locations are given in the Supplementary Table 1. The criteria for subject inclusion were: 1) presence of at least one hippocampal electrode with >100 SWRs recorded during the seizure-free overnight sleep; 2) presence of electrodes in at least one extrahippocampal region (amygdala, temporal or frontal cortices). The local field potential (LFP) was recorded during overnight sleep, typically starting 8:00-10:00 pm and lasting ~8-12 hours. The LFP was analog-filtered with 0.01 Hz highpass cutoff and recorded at 5000 Hz using the Nihon-Kohden recording system (256 channel amplifier, model JE120A) or at 8000 Hz using the Neuralynx ATLAS Clinical System. Sleep staging was performed in 30 sec blocks by a sleep specialist (B.A.M.), based on the visual inspection of scalp EEG at frontal, central, and occipital derivations, electrooculogram and electromyogram, guided by standard criteria^16,30^. The NREM sleep stages 2-4 (N2-4) were used in further analysis (Figure 1D).

### Electrode localization

Electrode localization was done using the pre-implantation MRI and post-implantation CT images. Both images were transformed into Talaraich space, followed by MRI segmentation (Freesurfer 5.3.067) and co-registration of T1-weighted structural MRI scans to the CT^51^. The electrode locations and selection of white matter contacts for re-referencing was verified by the epileptologist (J.J.L.).

### Recording locations

We analyzed SWRs recorded on 33 hippocampal locations (19 in left and 14 in right hemisphere; Figure 1B). The extra-hippocampal recording sites were grouped in three regions (Figure 1B): amygdala (including the basolateral, lateral and centromedial amygdala), temporal (including the insula, entorhinal, parahippocampal, inferior, medial and superior temporal cortices) and frontal cortex (including the orbitofrontal, medial prefrontal, dorsomedial and cingulate cortices). Regional distribution of extrahippocampal recording sites included: 36 (19 left, 17 right) in amygdala, 180 (106 left, 74 right) in temporal and 189 (89 left, 100 right) in frontal cortex (Figure 1C; Supplementary Table 1). Distribution of recording sites at individual subject, hemispheric and regional levels are shown in Table S1.

### Data Preprocessing

Recordings were re-referenced to the nearest white matter contact, resampled to 2000 Hz with linear interpolation (resample.m function in Matlab Signal Processing Toolbox) and high-pass filtered at 0.5 Hz using 4^th^ order Chebyshev filter. The data analysis and visualization was performed using the custom-written Matlab code, as well as the Freely Moving Animal (FMA; http://fmatoolbox.sourceforge.net/), Circular Statistics^52^ and FieldTrip^51,53^ toolboxes. Electrodes outside of the primary epileptogenic regions were used in the analysis. Further, interictal epileptic discharges (IEDs) on those electrodes were detected based on the combination of amplitude and derivative thresholds^20^. For amplitude-based IED detection, each LFP trace was low-pass filtered (300 Hz cutoff frequency) and the envelope of filtered trace was z-scored, while for the derivative-based IED detection, absolute differences between the consecutive voltage samples were z-scored. IEDs were detected based on the threshold crossing (mean + 5SD) by either the amplitude or derivative trace (Figure 1E). This method is optimized for detection of sharp transients that correspond to IEDs. Finally, automatic IED detection accuracy was validated by comparison with visual scoring performed by an epileptologist (Figure S1B).

### Sharp-wave ripple detection

Following electrode localization, LFP from hippocampal channels was bandpass-filtered in SWR range (80-150 Hz) using the 4^th^ order Chebyshev filter (filtfilt.m function in Matlab Signal Processing Toolbox), rectified and the upper envelope of rectified trace was z-scored. SWRs were detected using the FMA Toolbox, based on the double threshold crossing criteria: 1) envelope trace exceeding mean + 2 SDs for 20-100 ms and 2) the peak during this period exceeding mean + 5 SDs (Figure S1A). SWRs within 1 sec from nearest IED and hippocampal channels with <100 SWRs remaining after exclusion of SWRs in IED proximity were excluded from analysis.

### Peri-SWR HFA modulation

The HFA analytical amplitude was calculated by bandpass-filtering the raw LFP in 70-200 Hz range, Hilbert-transforming the filtered signal (hilbert.m function in Matlab Signal Processing Toolbox) and extracting the analytic signal amplitude. HFA amplitude trace was smoothed by convolving with Gaussian kernel (15 ms width, fastsmooth.m function in Matlab Signal Processing Toolbox) and binned (25 ms bin size), resulting in 20 time bins extending over the +/- 250 ms window centered at SWR peak (peri-SWR window). The time window between −500 and −250 ms relative to SWR peak was used as a baseline (Figure 2B). Baseline normalization was done at single trial level, by z-scoring concatenated baseline and peri-SWR HFA time windows. Peri-SWR HFA modulation was assessed at individual time bin level, by comparing the mean HFA within each bin with the mean HFA from the baseline period, using the two-tailed paired t-test (p < 0.05). Correction for multiple comparisons was done using Benjamini-Hochberg method^55^, based on the number of time bins within the peri-SWR window (n = 20). In addition, peri-SWR HFA modulation significance criteria at individual channel level included the presence of at least two consecutive time bins (a total of 50 msec) showing significant peri-SWR HFA modulation in the same direction. Target sites were first classified based on the presence (HFA+) or absence (HFA-) of significant peri-SWR HFA modulation. Based on the peri-SWR HFA modulation type, HFA+ sites were further classified as positively-, negatively- or mixed-modulated, the latter defined by the presence of both positive and negative modulation periods during peri-SWR window (Figure 2B). The time bin width choice (25 msec) was based on the fine temporal structure of peri-SWR cortical neuronal spiking fluctuations in rodents^6,42^ and humans^22^.

### Principal component analysis of peri-SWR HFA dynamics

Principal component analysis (PCA) of average peri-SWR HFA time-courses was performed on the pseudo-population consisting of HFA+ target sites from all the subjects, both ipsi- and contra-lateral to SWR location (n = 368). First, the peri-SWR HFA matrix (n x t) was constructed, with rows representing the individual target sitetrial-averaged and z-scored peri-SWR HFA time-courses (−250 to 250 ms), and columns representing 20 peri-SWR time bins (25 ms each). Covariance was calculated over all the target site pairs and the principal components were extracted by applying the eig.m Matlab function on peri-SWR HFA covariance matrix. Individual principal component (PC_n_) was considered significant if the percentage of explained variance (PC_nEV%)_ was at least 2-fold larger than the next PC (PC_n+1EV%_). The PC time-courses (Fig. 3B) were reconstructed by calculating the dot product between the given PC eigenvector and population vector activity at individual time bins. The regional PC time-courses were calculated using the PC weights from the target sites localized in a given region and calculating the dot product with the corresponding HFA population vector from the same region and time bin.Two-way analysis of variance (ANOVA) with region and hemisphere as the main factors was performed on the weight distribution of each significant PC, followed by planned comparisons using the two-tailed t-test (p < 0.05).

### SWR phase locking to target site slow-wave activity

The raw LFP recorded at each extra-hippocampal location was bandpass filtered in slow-wave range (0.5 - 4 Hz). Filtered traces were Hilbert-transformed and instantaneous phases were extracted using the angle.m function in Matlab. SWR-slow-wavephase locking is defined as a measure of consistency of local subcortical/cortical slow-wave phase at the times of hippocampal SWR peaks (Figure 4A). For each target site, significance of SWR phase locking to slow-wave in target site was quantified using the Rayleigh test (circ_rtest.m function, Matlab Circular Statistics Toolbox^52^). Multiple comparisons correction was performed using the Benjamini-Hochberg method^55^, based on the number of recording sites in a given subject. SWR-slow-wave synchrony was defined based on presence or absence of significant SWR-slow-wave phase locking (SWR-slow-wave+ or SWR-slow-wave-). For each target site showing SWR-slow-wave synchrony (SWR-slow-wave+), mean SWR-slow-wave phase angle was computed using circ_mean.m function (Matlab Circular Statistics Toolbox). Peri-SWR HFA modulation strength on a given target site (Figure 4C) was defined as the summed absolute values of a z-scored HFA trace within a 100 msec sliding window (25 msec step size), with centers starting −200 msec and ending 200 msec following SWR. SWR-slow-wave synchrony magnitude was defined as the length of vector r, an output of Rayleigh test (circ_rtest function from Matlab Circular Statistics Toolbox). Vector r could take any value in the range 0-1 and the higher value denotes SWRs occurring more consistently at given phase of target site slow-wave. For each target site, correlation between the SWR-slow-wave synchrony magnitude and peri-SWR HFA modulation strength was calculated using Spearman correlation (Figure 4C).

To verify that SWR-slow-wave phase locking is not driven by higher slow-wave amplitude on a given target site, LFP signal from each target site was filtered in slow-wave range (0.5 - 4 Hz) and the target site slow-wave average amplitude was computed as the mean of slow-wave analytical amplitude from individual peri-SWR windows.Comparison of peri-SWR slow-wave amplitude between theSWR-slow-wave+ and SWR-slow-wave- target sites was done using the two-tailed t-test for independent samples (p < 0.05; Figure S3D). The lack of significant group difference in peri-SWR slow-wave amplitude was interpreted as ruling out the possibility of peri-SWR slow-wave amplitude having a confounding effect on slow-wave phase extraction and SWR-slow-wave phase locking calculation.

### Generalized linear model

The generalized linear model (GLM) analysis was performed by modelling the average peri-SWR HFA time-course on a target site A (HFA-A) as a function of HFA time-course of another simultaneously recorded target site (HFA-B) and averaged population peri-SWR HFA time-course (HFA-pop), with population defined as all the simultaneously recorded target sites, except A and B. All of the analysis was done at zero time lag. To prevent the influence ofabsolute HFA levels on the resulting coupling estimates, individual peri-SWR HFA time-courses were z-scored. The model fitting was done using a Matlab glmfit.m function, with normal distribution. For each target site pair, the model produced two coefficients (β_peer_ and β_pop_), reflecting the predictive power of HFA-B and HFA-pop on HFA-A. β_peer_ at zero time lag (β_0_) was used in further analysis. Intra-modular target sites were defined based on the presence of both significant peri-SWR HFA modulation (HFA+) and SWR-slow-wave synchrony. For each simultaneously recordedintra-modular target site pair (A and B) with mean SWR-slow-wave phase angles Φ_A_ and Φ_B_, the absolute SWR-slow-wave phase difference (Φ_diff_)was computed as follows:

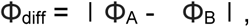

with Φ_a_ and Φ_b_ representing the mean phase angles on the respective target sites, across all the SWRs recorded on a given hippocampal channel. If the individual within- or cross-regional β_0_-Φ_diff_ correlation was significant (Spearman correlation, p<0.05), we compared the distributions of β_0_ coefficients between the intra-modular and extra-modular target site pairs, the latter defined as the pairs containing just intra-modular target site. Distributions of β_0_ between each categorywere compared using Ansari-Bradley tests (p<0.05). The variance of intra- and extra-modular β_0_was expressed as the standard error of mean (SEM) and calculatedforsliding windows (100 msec width, 25 msec step size), centered between −200 and 200 msec around SWR peak. As the unequal sample size could affect the variance calculation, we randomly sampled 100 sets of extra-modular target site pairs, equal in size to the intra-modular set (n = 166). Significant difference was reached if the intra-modular variance during given time window exceeded 99th percentile of the extra-modular set (p < 0.01; Figure 5E).

### Statistical Analysis

The analysis was done using all the hippocampal channels that passed the inclusion criteria. This approach was chosen because the SWRs at different hippocampal locations might be associated with different dynamics in the same extra-hippocampal populations^54^. In addition, including the hippocampal channels from both hemispheres allowed the testing of both ipsi- and contralateral brain dynamics during peri-SWR periods. Therefore, the peri-SWR dynamics in all the extra-hippocampal recording sites was analyzed relative to SWRs on each individual different hippocampal site in the same subject. For the purpose of this analytical approach, each extra-hippocampal recording site was defined as a target site, with respect to individual hippocampal site. Target sites in each region were classified as ipsi- or contralateral relative to SWR location. The regional distribution of target sites included: 87 (44 ipsilateral, 43 contralateral) in the amygdala, 521 (250, 271) in temporal and 700 (331, 369) in frontal cortex.

All the corrections for multiple comparisons were done using the Benjamini-Hochberg method^55^. Correction factor was based on the number of time bins during peri-SWR window for individual target site peri-SWR HFA modulation, the number of recording locations in individual patient for the SWR-slow-wave phase locking or the number of regional and hemispheric comparisons in other cases. Analysis was performed on the pseudo-populations, containing all the target sites that passed inclusion criteria for a given analysis. The target sites with at least 50 IED-free peri-SWR windows (± 1 sec) were used for peri-SWR HFA modulation (Figures 2, S2), SWR-slow-wave phase locking (Figure 4, S3) and GLM-based HFA pairwise coupling analysis (Figure 5; n = 1308). PCA included all the target sites showing significant peri-SWR HFA modulation (Figure 3; n = 368). Correlation between the target site pairwise SWR-slow-wave locking phase difference (φ_diff_)and HFA coupling coefficient (β_0_) was done for target sites showing both significant peri-SWR HFA modulation and significant SWR-slow-wave phase locking (intra-modular; n = 234). Main effects of region and hemisphere, as well as region*hemisphere interactions in pseudo-populations were obtained using the linear regression (Figures 2, S2, 3), followed by planned comparisons using chi-square test (p < 0.05). Uniformity of SWR-slow-wave phase distributions was tested using the Rayleigh test (Figure 4; p < 0.05).

## General

We wish to thank Aaron Wilber for the useful comments and discussion, as well as the patients, nurses, technicians, and physicians at the University of California Irvine Epilepsy Unit.

## Author contributions

Conceptualization, I.S. and J.J.L.; Methodology, I.S., J.Z., H.Z., B.A.M., S. M., O.K.M., S.V.; Surgeries, S.V.; Investigation, I.S., J.Z.; Formal Analysis, I.S.; Writing - Original Draft, I.S.; Writing - Review & Editing, I.S. and J.J.L., with input from other authors; Funding Acquisition, I.S., B.L.M., R.T.K. and J.J.L.; Resources, B.L.M., R.T.K. and J.J.L.; Supervision, J.J.L.

## Data and materials availability

Data and code are available from the corresponding author upon reasonable request.

## Supplementary Materials

**Supplementary Fig.1.**
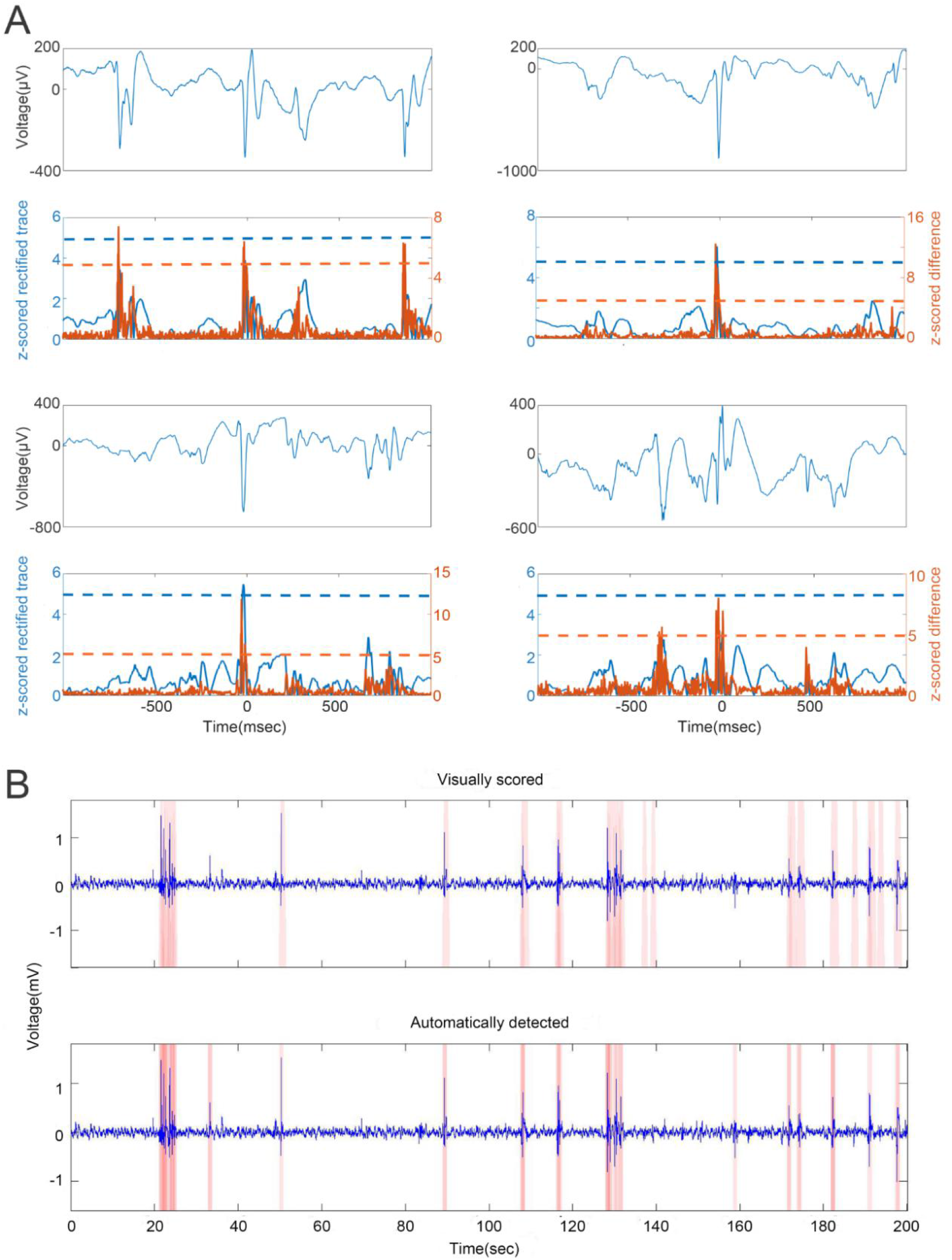
Detection of interictal epileptic discharges (IEDs). a, Examples of detected interictal epileptic discharges (IEDs) and illustration of IED detection algorithm. The top plot in each example: raw LFP trace around the IED. Bottom plot in each example: Blue - z-scored rectified raw trace. Orange - z-scored derivative of raw trace. Detection was based on any of the two traces crossing the threshold of mean + 5SD (dashed lines). The periods of ± 1 sec around the detected IEDs were excluded from the analysis. b, Example of the raw local field potential (LFP) trace with IEDs (pink bars) annotated by a trained observer (top). The same LFP trace with IEDs detected by an automatic algorithm (pink bars, bottom).

**Supplementary Fig.2.**
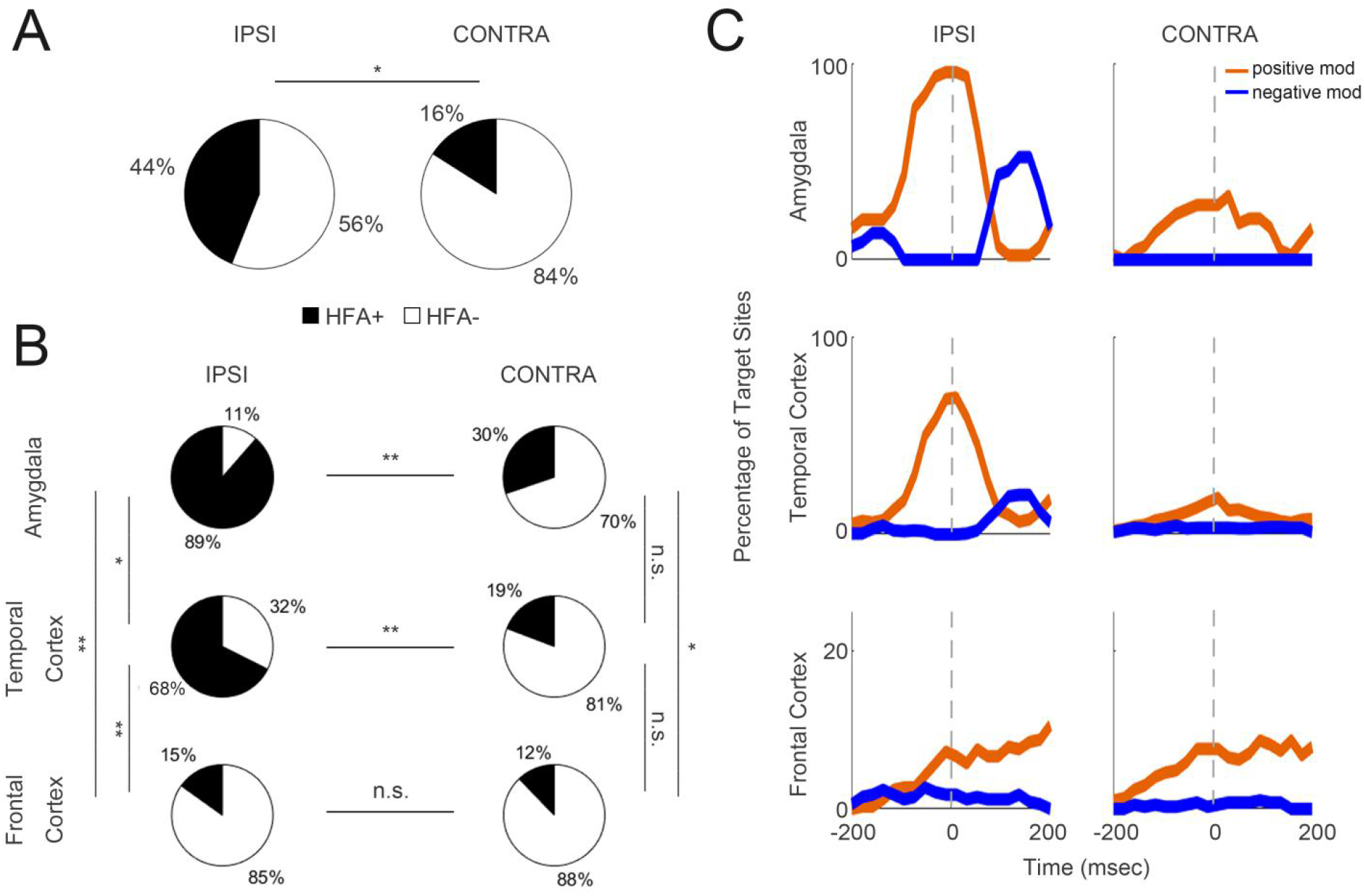
Regional and hemispheric distributions of peri-SWR HFA modulation. **a,** Percentage of significant high frequency activity modulation (HFA+) around SWRs is higher in the hemisphere ipsilateral to SWR location, relative to contralateral (χ2 (1, n = 1308) = 116.2, p < 10^-10^). Black: HFA+. White: HFA-. **b**, Percentages of HFA+ are significantly higher in the amygdala and temporal cortex, relative to frontal cortex. HFA+ percentages are higher ipsilateral to SWR location in amygdala and temporal cortex, but not in the frontal cortex (χ2 test, p < 0.05, statistical details in Supplementary Table 2B). Black: HFA+. White: HFA-. **c**, Regional percentages of target sites showing significantlypositive (orange) or negative (blue) HFA modulation at a given time bin during the peri-SWR window. Left column - ipsilateral and right column - contralateral to SWR location. SWR peak time is shown as dashed line. Black: HFA+. White: HFA-.

**Supplementary Fig.3.**
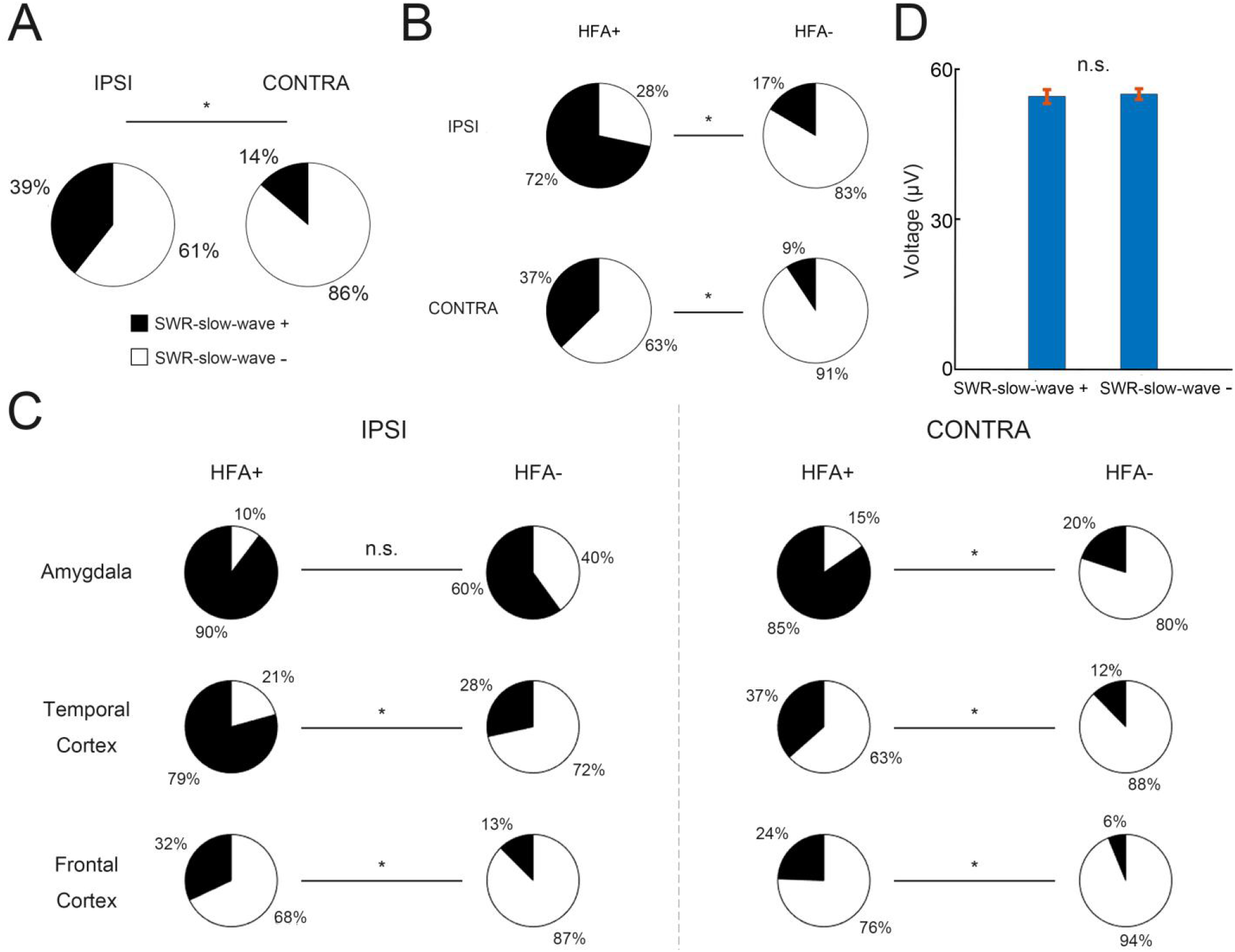
SWR synchrony with subcortical/cortical slow-waves (SWR-slow-wave +) predicts local peri-SWR HFA modulation at the hemispheric and regional level. **a,** Percentage of SWR-slow-wave+ target sites is significantly higher ipsilateral to SWR location. Pseudo-population consisting of 625 ipsilateral and 683 contralateral locations (χ2(1, n =1308) = 131.3, p < 10^-10^). Black: SWR-slow-wave+. White: SWR-slow-wave-. **b**, Right: Percentages of SWR-slow wave+ phase locking are higher for the target sites with significant peri-SWR HFA modulation (HFA+), relative to target sites without significant peri-SWR HFA modulation (HFA-), both ipsilateral (χ2 (1, n =625) = 182.5, p < 10^-10^) and contralateral (χ2 (1, n =683) = 53.4, p < 10^-10^) to SWR location. Black: SWR-slow-wave+. White: SWR-slow-wave-.**c**, Percentages of SWR-slow wave+ phase locking are significantly higher for the target sites with significant peri-SWR HFA modulation (HFA+) in all the regions ipsilateral and contralateral to SWR location (χ2> 10, all p’s < 10^-3^), except the ipsilateral amygdala (χ2(1, n =44) = 3.3, p = 0.07). Black: SWR-slow-wave+. White: SWR-slow-wave-. **d**, Slow-wave amplitude during peri-SWR period (± 1 sec) does not differ between the SWR-slow wave+ and SWR-slow wave- target sites (t(1306) = 0.19, p = 0.85; n = 1308). The data is shown as mean ± SEM. Black: SWR-slow-wave+. White: SWR-slow-wave-.

**Supplementary Table 1.** Recording site distributions across the subjects, hemispheres and regions. L = Left, R = Right. HIPP = hippocampus, AMY = amygdala, TEMP = temporal cortex, FRON = frontal cortex.

| Subject ID | Sex, Age | HIPP (L, R) | AMY (L, R) | TEMP (L, R) | FRON (L, R) |
| --- | --- | --- | --- | --- | --- |
| 1 | M, 41 | 1, 0 | 1, 3 | 5, 11 | 16, 11 |
| 2 | F, 32 | 1, 0 | 4, 4 | 12, 5 | 3, 12 |
| 3 | F, 48 | 2, 1 | 0, 3 | 4, 10 | 9, 9 |
| 4 | M, 24 | 4, 2 | 3, 0 | 8, 13 | 0, 21 |
| 5 | F, 55 | 1, 2 | 0, 2 | 13, 3 | 2, 9 |
| 6 | F, 33 | 1, 0 | 0, 0 | 4, 0 | 0, 0 |
| 7 | M, 56 | 1, 0 | 1, 0 | 9, 0 | 0, 0 |
| 8 | M, 28 | 4, 0 | 2, 2 | 5, 6 | 11, 9 |
| 9 | M, 27 | 1, 2 | 0, 0 | 14, 7 | 13, 9 |
| 10 | M, 57 | 1, 0 | 3, 0 | 10, 0 | 7, 0 |
| 11 | F, 25 | 2, 4 | 2, 2 | 8, 12 | 28, 20 |
| 12 | M, 29 | 0, 3 | 3, 1 | 14, 7 | 0, 0 |

**Supplementary Table 2A.** The numbers and percentages of target sites showing significant peri-SWR high frequency activity modulation in each region and hemisphere (ipsi- or contralateral to SWR location). Ipsilateral and contralateral denotes the target site hemisphere, relative to SWR location.

| Ipsilateral | Peri-SWR HFA modulation |
| --- | --- |
| Amygdala | 39/44 (88.6%) |
| Temporal | 189/250 (67.6%) |
| Frontal | 50/331 (15.1%) |
| Contralateral | Peri-SWR HFA modulation |
| Amygdala | 13/43 (30.2%) |
| Temporal | 52/271 (19.2%) |
| Frontal | 45/369 (12.2%) |

**Supplementary Table 2B.** Chi-square statistics for the regional and hemispheric percentages of peri-SWR HFA modulation (p < 0.05, Benjamini-Hochberg correction for multiple comparisons; *p<0.005,**p<0.0005). Ipsilateral and contralateral denotes the target site hemisphere, relative to SWR location.

| **Ipsilateral** | **Chi-square statistics** |
| --- | --- |
| **Temporal-Frontal\*\*** | $\chi^2(1,580) = 167.1; p < 10^{-10}$ |
| **Temporal-Amygdala\*** | $\chi^2(1,293) = 8.0; p < 0.005$ |
| **Frontal-Amygdala\*\*** | $\chi^2(1,374) = 116.0; p < 10^{-10}$ |
| **Contralateral** | **Chi-square statistics** |
| **Temporal-Frontal** | $\chi^2(1,639) = 5.9; p < 0.05$ |
| **Temporal-Amygdala** | $\chi^2(1,313) = 2.8; p = 0.10$ |
| **Frontal-Amygdala\*** | $\chi^2(1,411) = 10.4; p < 0.005$ |
| **Interhemispheric** | **Chi-square statistics** |
| **Temporal-Temporal\*\*** | $\chi^2(1,520) = 124.8; p < 10^{-10}$ |
| **Frontal-Frontal** | $\chi^2(1,699) = 1.3; p = 0.26$ |
| **Amygdala-Amygdala\*\*** | $\chi^2(1,86) = 30.8; p < 10^{-7}$ |

**Supplementary Table 3.** T-test statistics for the regional comparisons of the first three principal component weights (p < 0.05, Benjamini-Hochberg correction for multiple comparisons; *p<0.005,**p<0.0005). Ipsilateral denotes the target site hemisphere, relative to SWR location.

|  | **PC1** | **PC2** | **PC3** |
| --- | --- | --- | --- |
| **Region** | $F(2,367) = 133.1$<br>**$p < 10^{-10}$ | $F(2,367) = 37.4$<br>**$p < 10^{-10}$ | $F(2,367) = 7.3$<br>*$p < 0.001$ |
| **Hemisphere** | $F(1,367) = 17.8$<br>*$p < 10^{-4}$ | $F(1,367) = 8.7$<br>*$p < 0.005$ | $F(1,367) = 15.0$<br>*$p < 0.005$ |
| **Region Hemisphere Interaction** | $F(2,367) = 32.4$<br>**$p < 10^{-10}$ | $F(2,367) = 11.6$<br>**$p < 10^{-10}$ | $F(2,367) = 2.4$<br>$p = 0.1$ |
| **Temporal - Frontal** | $t(1,217) = -18.6$<br>**$p < 10^{-10}$ | $t(1,217) = 10.5$<br>**$p < 10^{-10}$ | $t(1,217) = -3.4$<br>*$p < 0.005$ |
| **Temporal - Amygdala** | $t(1,206) = 2.86,$<br>$p = 0.01$ | $t(1,206) = 3.83,$<br>*$p < 10^{-3}$ | $t(1,206) = 1.53,$<br>$p = 0.11$ |
| **Frontal - Amygdala** | $t(1,87) = 15.6,$<br>*$p < 10^{-10}$ | $t(1,87) = 8.29,$<br>**$p < 10^{-10}$ | $t(1,87) = 1.43,$<br>$p = 0.16$ |

**Supplementary Table 4.** Chi-square statistics for the comparisons of SWR phase locking to slow waves on target sites showing significant or non-significant peri-SWR high frequency activity modulation (p < 0.05, Benjamini-Hochberg correction for multiple comparisons; *p<0.005,**p<0.0005). Ipsilateral and contralateral denotes the target site hemisphere, relative to SWR location.

| Ipsilateral | HFA + sites | HFA- sites | Chi-square statistics |
| --- | --- | --- | --- |
| Temporal** | 79.3% (134/169) | 28.4% (23/81) | $\chi^2(1,250) = 60.7, p < 10^{-10}$ |
| Frontal* | 32.0% (16/50) | 12.5% (35/281) | $\chi^2(1,331) = 12.4, p < 10^{-3}$ |
| Amygdala | 89.7% (35/39) | 60.0% (3/5) | $\chi^2(1,44) = 3.3, p = 0.07$ |
| Contralateral | HFA + sites | HFA- sites | Chi-square statistics |
| Temporal* | 36.5% (19/52) | 12.3% (27/219) | $\chi^2(1,271) = 17.5, p < 10^{-4}$ |
| Frontal* | 24.4% (11/45) | 6.2% (20/324) | $\chi^2(1,370) = 17.1, p < 10^{-4}$ |
| Amygdala* | 84.6% (11/13)<sup>2</sup> | 20.0% (6/30) | $\chi^2(1, 43) = 15.8, p < 10^{-4}$ |

**Supplementary Table 5.** Correlation between the beta coefficient and phase difference at the whole brain, within- and cross-regional levels. Only the region pairs with more than 20 target site pairs were included in the analysis (Spearman correlation; *p < 0.05; Benjamini-Hochberg correction for multiple comparisons).

|  |  |
| --- | --- |
| All pairs | $r = -0.04; p = 0.05; n = 1716$ pairs |
| Ipsilateral |  |
| Temporal-Frontal* | $r = -0.26; p < 0.001; n = 166$ pairs |
| Temporal-Amygdala | $r = 0.10; p = 0.09; n = 302$ pairs |
| Frontal-Amygdala | $r = 0.10; p = 0.60; n = 32$ pairs |
| Temporal-Temporal | $r = -0.09; p = 0.04; n = 450$ pairs |
| Contralateral |  |
| Temporal-Temporal | $r = -0.41; p = 0.02; n = 32$ pairs |
| Inter-Hemispheric |  |
| Temporal-Temporal | $r = -0.02; p = 0.75; n = 176$ pairs |

**Movie S1.Temporal dynamics of significant peri-SWR HFA modulations in the hemisphere ipsilateral (left) or contralateral (right) to SWR location.** The time window includes ±250 ms around the SWR peak, binned into 25 msec time bins (20-time bins in total). Significant HFA modulations were more prevalent in the hemisphere ipsilateral to SWR location and in the temporal lobe (amygdala and temporal cortex), relative to the frontal cortex. Orange or blue markers denote the anatomical locations of significant positive or negative modulations at a given time during the peri-SWR window. Note the widespread HFA decrease ~100 msec following the SWR peak in the ipsilateral temporal lobe, but not in the contralateral hemisphere.

